# Where in the brain do internally generated and externally presented visual information interact?

**DOI:** 10.1101/2023.07.10.548319

**Authors:** Jussi Alho, Athanasios Gotsopoulos, Juha Silvanto

**Author notes:** Corresponding author: Jussi Alho, P.O. Box 21, Haartmaninkatu 3, Helsinki, FI-00014, Finland. These authors contributed equally (Jussi Alho, Athanasios Gotsopoulos).

## Abstract

Conscious experiences normally result from the flow of external input into our sensory systems. However, we can also create conscious percepts independently of sensory stimulation. These internally generated percepts are referred to as mental images, and they have many similarities with real visual percepts. Consequently, mental imagery is often referred to as “seeing in the mind’s eye”. While the neural basis of imagery has been widely studied, the interaction between internal and external sources of visual information has received little interest. Here we examined this question by using fMRI to record brain activity of healthy human volunteers while they were performing visual imagery that was distracted with a concurrent presentation of a visual stimulus. Multivariate pattern analysis (MVPA) was used to identify the brain basis of this interaction. Visual imagery was reflected in several brain areas in ventral temporal, lateral occipitotemporal, and posterior frontal cortices, with a left-hemisphere dominance. The key finding was that imagery content representations in the left lateral occipitotemporal cortex were disrupted when a visual distractor was presented during imagery. Our results thus demonstrate that the representations of internal and external visual information interact in brain areas associated with the encoding of visual objects and shapes.

## 1 Introduction

Visual imagery allows us to internally generate mental representations of visual objects independently of external visual stimulation (e.g., Kosslyn, 2005; Pearson, Naselaris, Holmes, & Kosslyn, 2015). Studies examining the blood-oxygen-level dependent (BOLD) activation with functional magnetic resonance imaging (fMRI) have found that imagery and perception of visual objects evoke content-related activation in the same occipitotemporal category-selective brain areas (Ishai, Ungerleider, & Haxby, 2000; O’Craven & Kanwisher, 2000). However, BOLD-fMRI activation in the same brain area does not conclusively disclose whether the neural representations are shared. Advances in fMRI data analysis brought by machine-learning-based multivariate pattern analysis (MVPA) methods allow a more scrupulous inspection of neural representations. MVPA differs from conventional univariate fMRI analyses in that it considers whether information related to a specific stimulus or experimental condition is encoded in activity patterns across multiple voxels. In MVPA, a classifier can be trained to identify BOLD-fMRI activity patterns related to a specific stimulus or experimental paradigm. If the trained classifier can then classify unseen data to the correct stimulus or condition categories, a conclusion can be made that the stimulus or mental operation required in the paradigm is represented in the brain area(s) of the learned activation pattern.

Prior studies using MVPA on fMRI data have revealed shared neural representations of imagined and perceived objects in visual brain areas, especially within the ventral visual processing stream (Radoslaw M. Cichy, Heinzle, & Haynes, 2012; Lee, Kravitz, & Baker, 2012; Reddy, Tsuchiya, & Serre, 2010). However, brain lesion studies have showed a double dissociation between imagery and perception. For example, a patient with damage in the ventral pathway was unable to recognize objects yet had intact ability to form visual images (Behrmann, Moscovitch, & Winocur, 1994), whereas another patient with a lesion in the associative visual pathways (i.e., the secondary visual cortex and both dorsal and ventral visual streams) exhibited impaired visual imagery yet was able to recognize visually presented objects (Farah, Levine, & Calvanio, 1988). In sum, it seems that while many neural mechanisms between visual perception and imagery are shared, there are also evident differences. Indeed, the lack of bottom-up input from the retina must be somehow compensated in visual imagery.

While numerous studies have investigated the neural basis of both visual imagery and perception, the interaction between imagery and external visual input has received little interest. This despite various interesting phenomena indicating a loose link between the two. In a seminal experiment, Perky (1910) asked participants to visualize various objects on a screen while, unbeknownst to the participants, a faint real image matching the imagined object was projected onto the screen. When asked afterwards, the participants believed that the projected real images were a product of their imagination. Such similarity of the subjective experience implies similarity also in the neural representations between perception and imagery of visual objects.

Although vision-imagery interaction has been understudied, interaction of vision with visual short-term memory (VSTM), known to be closely related to visual imagery (e.g., Albers, Kok, Toni, Dijkerman, & de Lange, 2013), has received more attention. Using MVPA measures on fMRI data, recent work found that distracting visual information affects representations of VSTM in occipital cortex but not in parietal cortex (Bettencourt & Xu, 2016; Lorenc, Sreenivasan, Nee, Vandenbroucke, & D’Esposito, 2018), thus identifying the locus of vision-VSTM interaction to visual cortical areas.

Our aim was to examine where in the brain does the interaction between perception and imagery of visual objects occur. This was accomplished by using a paradigm in which visual distractors were presented while participants were engaged in visual imagery. We measured BOLD-fMRI responses from eighteen participants who were performing a task involving imagery of visual object shapes. In half of the imagery trials a low-visibility visual distractor was presented during the imagery period. A linear artificial neural network classifier was trained on the fMRI activation patterns to differentiate between trial types. To identify the brain area(s) most contributing to the classification, we used state-of-the-art importance extraction methods. In line with findings from VSTM, we hypothesized that interaction between imagery and perception occurs in cortical areas where the incoming visual input is encoded, rather than in higher-level (e.g., parietal or frontal) areas associated with conscious experience.

## 2. Materials and methods

### 2.1 Participants

Twenty-one volunteers participated in the experiment. Three participants were excluded from the analyses due to inadequate behavioral performance (see section 3.1), resulting in final sample of eighteen participants (ages 19–49 years, 9 males, mean age 31.2 years). All participants were healthy with normal or corrected-to-normal vision. Written informed consent was obtained from each participant. The study protocol was approved by the Ethics Committee of the Hospital District of Helsinki and Uusimaa.

### 2.2 Experimental Design

The experimental paradigm (adapted from Jacobs, Schwarzkopf, & Silvanto, 2018) is depicted in **Figure 1**. Participants were first shown one of three letters (Y, R, or E) located in one of the corners of the screen. In case of letter Y (the Finnish initial for *circle*) or R (the Finnish initial for *diamond*, i.e., a square standing on its corner), the participants were to imagine a circle or a diamond, respectively, inside the placeholders for the duration that the placeholders were visible on the screen (4000 ms). The participants were instructed to imagine as large a shape as they can fit inside the placeholders. In case of letter E (the Finnish initial for *nothing*), participants were not required to imagine anything, but merely to fixate on the centre of the placeholders (null trial). In half of the trials, a low-visibility visual distractor (*triangle*) was presented for 20 ms, with randomly varying onset of 200-500 ms from the onset of the imagery period (i.e., placeholders appearing on the screen). Thus, the experimental design involved altogether six trial types: 1) *circle imagery without distractor*, 2) *diamond imagery without distractor*, 3) *no imagery without distractor*, 4) *circle imagery with distractor*, 5) *diamond imagery with distractor*, 6) *no imagery with distractor*.

**Figure 1:**
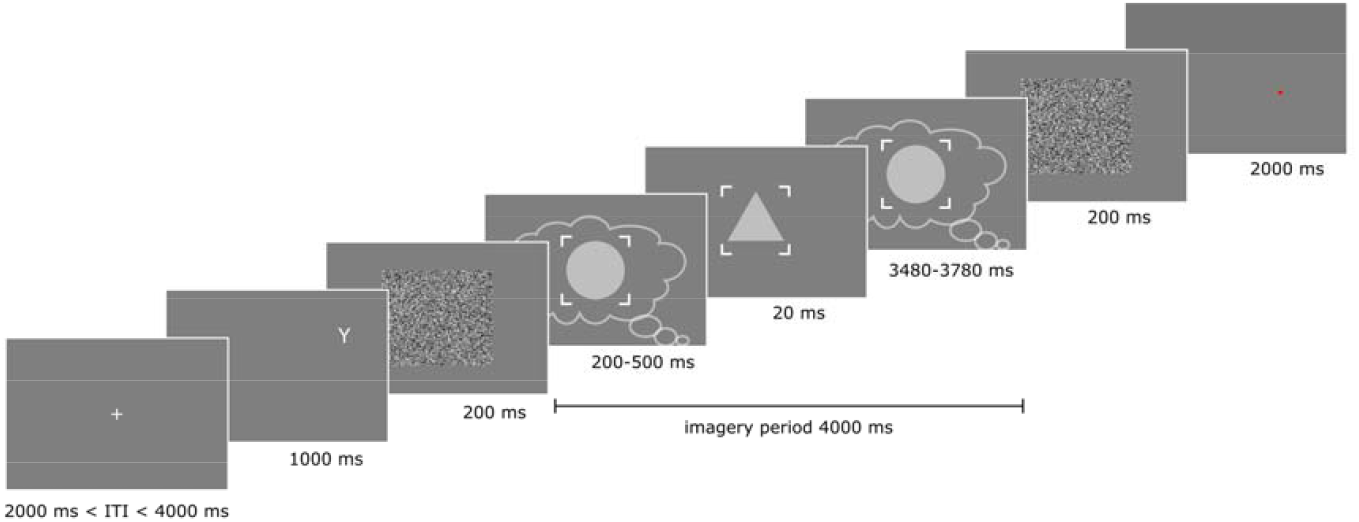
Experimental paradigm. example of a trial where circle was to be imagined and distractor was presented. After the inter-trial interval, a letter was shown indicating the shape to be imagined. The letter was followed by a visual noise mask after which placeholders appeared on the screen to indicate the area within which the shape was to be imagined. In half of the trials, low-visibility visual distractor (triangle) was presented during the imagery period. After the place holders disappeared (signifying the end of the imagery period), another visual noise mask appeared followed by a presentation of a red dot in a random location on the screen. The participants were to indicate with a button press whether the red dot fell inside the shape they had imagined.

After the imagery period, a red dot appeared on the screen near the edges of the imagined shape and the participants were to indicate with a button press whether the dot fell inside its area. The purpose of this task was to keep the participants alert, to control for their level of attention, and to investigate the effect of the visual distractor on behaviour. Both the instruction letter and the imagery period were followed by a 200-ms visual white noise mask to reduce after-image effect. Each trial was followed by a 2000-4000 ms inter-trial interval. The experiment comprised six runs, each consisting of 48 trials (16 for each trial type), resulting in a total scanning time of approximately one hour per participant. The ordering of the trial types within a run was fully randomized to avoid any ordering effects (due to the possibility of the response in the previous trial still being present in the signal).

The stimuli were delivered using Presentation software (Neurobehavioral Systems, Inc., Albany, CA, USA) and back-projected to the participant via a mirror and a semitransparent screen using a 3-micromirror data projector (Christie X3, Christie Digital Systems Ltd, Mönchengladbach, Germany).

### 2.3 fMRI Data Acquisition and Preprocessing

MRI data were collected on a 3T Siemens Magnetom Skyra scanner at the Advanced Magnetic Imaging Centre, Aalto NeuroImaging, Aalto University, using a 32-channel Siemens volume coil. Whole-brain functional scans were collected using a whole-brain T2*-weighted EPI sequence, sensitive to the blood-oxygen-level dependent (BOLD) contrast, with the following parameters: 33 axial slices, slice thickness 4 mm, matrix size = 64 × 64, echo time (TE) = 24 ms, repetition time (TR) = 1.7 s, flip angle = 75°, and field-of-view (FOV) = 20 cm. High-resolution anatomical images with isotropic 1 × 1 × 1 mm^3^ voxel size were collected using a T1-weighted MP-RAGE sequence.

The fMRI data were preprocessed using the FSL software (www.fmrib.ox.ac.uk) and custom MATLAB code (BRAMILA pipeline v2.0, available at https://version.aalto.fi/gitlab/BML/bramila). EPI slices were first corrected for slice timing differences. Volumes were then corrected for head motion using MCFLIRT and coregistered with FLIRT to the Montreal Neurological Institute (MNI) 152 2mm template in a two-step registration procedure: first from EPI to the individual anatomical image after brain extraction (9 degrees of freedom) and then from the anatomical image to standard template (12 degrees of freedom). To remove scanner drift, a high pass temporal filter (cut-off frequency of 0.01 Hz) was applied to the time series. To further control for motion and physiological artefacts, BOLD time series were cleaned using 24 motion-related regressors, signal from deep white matter, ventricles and cerebral spinal fluid locations as described in Power et al 2012.

### 2.4 Multivariate Pattern Analysis (MVPA)

The motivation to analyze our data using a neural network classifier, instead of univariate analyses, such as GLM, is twofold: First, classification accuracies directly quantify the discriminability across conditions. Second, previous work has shown that importance maps generated from neural network classifiers can reveal multivariate patterns and patterns with low univariate information (Gotsopoulos et al., 2018). Such information may not be available when performing univariate analysis, such as GLM, or when feeding the classifier with data preprocessed by a univariate method, such as GLM coefficients.

Given the timing of the visual distractor (see section 2.2) and accounting for the hemodynamic lag (∼5 s), the fourth fMRI time point (5.1-6.8 s) after the imagery onset was input to the classifier. As significant difference was observed in the hit rates between trials with and without distractor (see section 3.1), we regressed out the subjective difficulty of the task (defined as the difference in the hit rates between trials with and without distractor) and z-scored (transformed to a mean of zero and unit variance) the fMRI data before the classification. A group brain mask was determined by selecting voxels that were present (i.e., had non-zero standard deviation values) in all participants, resulting in a total of 203477 voxels. Classification of different trial types was performed with a linear (i.e., no hidden layers) artificial neural network classifier, as implemented in an in-house developed neural network toolbox that has been previously used to classify fMRI data (Gotsopoulos et al., 2018; available at https://github.com/gostopa1/DeepNNs). The classifier utilized softmax activation function in the output layer and cross-entropy loss function. Training was performed for 5000 epochs using stochastic gradient descent as an optimization algorithm. Learning rate was set to 0.001. A low minibatch size of 20 was selected to avoid overfitting (Keskar, Mudigere, Nocedal, Smelyanskiy, & Tang, 2016). In the case of our linear neural network classifier, the output for each category was a normalized weighted sum of the inputs. The normalization occurs due to the softmax activation function that causes the output values to sum to 1. The input to output weights are updated in each training iteration based on the derivative of the loss function with respect to the weights of the neural network, that is, the essence of the back-propagation algorithm.

We used a within-participants classification approach with a leave-one-out cross validation where the classifier was trained with n-1 measurement runs and tested with the remaining run. Thus, a total of 40 samples (i.e., 5 runs x 8 trials) per trial type were available in training and 8 (1 run x 8 trials) samples per trial type in testing. Cross-validation was performed across all runs and the classification accuracy was calculated as an average percentage of correct predictions across the cross-validation runs. Feature selection was performed on the training data for each classification run using one-way ANOVA across the categories to exclude voxels whose activation was not modulated (i.e., showed no statistically significant differences between the means of the samples belonging to the different categories at p < 0.05). Statistical significance of the group-level classification accuracies was tested by generating a null-distribution by randomly shuffling the output labels of the whole data set before running the classifier. This process was repeated 1000 times. The p-value was calculated as the proportion of permutation runs where the value was greater than or equal to the observed value.

To determine the voxels most contributed to the classifier’s selection of a category, we extracted the importance values for each voxel using layer-wise relevance propagation, a method that distributes the output of the classifier to the inputs based on the weight-input product as suggested by previous work (Bach et al., 2015; Montavon, Lapuschkin, Binder, Samek, & Müller, 2017). The sum of the importance values is equal to the output of the classifier, thus providing a direct interpretation of each importance map (see Figure 2). The inputs and outputs (i.e., predicted values for the test set) were used for generating the importance maps, in order to minimize any overfitting-related effects. Participant-wise importance maps were calculated by taking the average over the cross-validation runs. For the group-level importance maps, the participant-wise importance maps were averaged across all participants for each category. **Figure 2** depicts the classification workflow.

**Figure 2:**
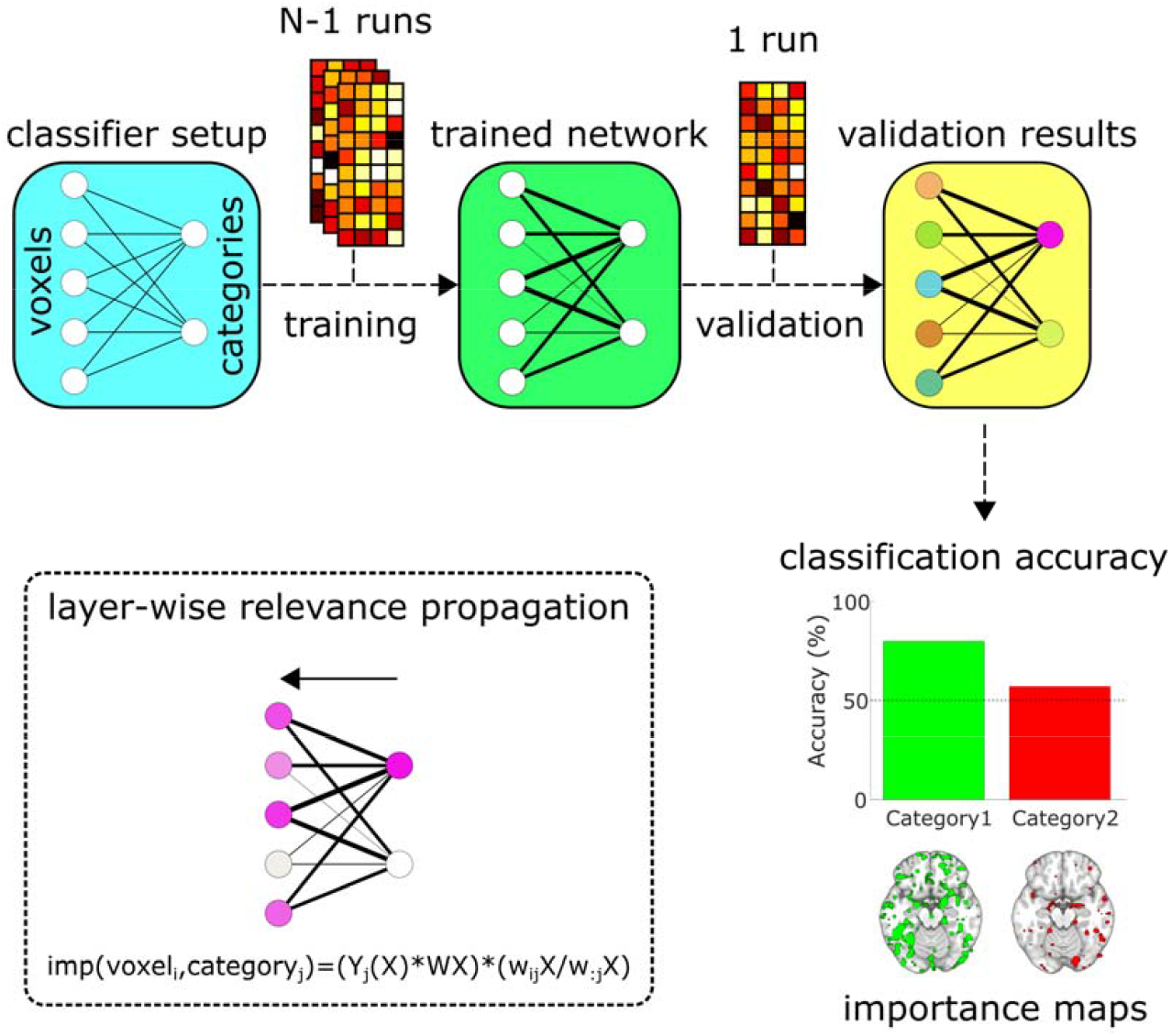
Classification workflow. An illustration of the steps of the MVPA together with a visual representation of the layer-wise relevance propagation method used in the importance extraction. Importance maps reveal the brain regions that contribute most significantly to the classification. Imp=importance; W = weights of the trained classifier; X = voxel activation; Y = classifier output. Modified from Gotsopoulos et al. (2018).

As a first analysis, we classified the *no imagery without distractor* and *no imagery with distractor* trials. In case of significantly above chance level classification accuracies, we can disclose the cortical regions representing the visual distractor by determining the voxels that most contributed (i.e., the highest importance values) to the classifier’s selection of the *no imagery with distractor* category. For the main analysis, we used both within- and cross-classifications. That is, we first trained a classifier to differentiate the *circle imagery without distractor* and *diamond imagery without distractor* trials. This classifier was then tested on both the *circle imagery without distractor* and *diamond imagery without distractor* trials (within-classification) as well as on the *circle imagery with distractor* and *diamond imagery with distractor* trials (cross-classification). Significantly lower classification accuracies in testing on the *circle imagery with distractor* vs. *diamond imagery with distractor* trials compared to the *circle imagery without distractor* vs. *diamond imagery without distractor* trials implies that the distractor interferes with the imagery content representations. However, comparison of within- and cross-classification might not be sufficient to show a reduction of information as cross-classification is almost always likely to have lower classification accuracy compared to within-classification. For this reason, we also trained a classifier to differentiate the *circle imagery with distractor* and *diamond imagery with distractor* trials. We tested this classifier on both the *circle imagery with distractor* and *diamond imagery with distractor* trials (within-classification) as well as on the *circle imagery without distractor* and *diamond imagery without distractor* trials (cross-classification). If information about the imagery content is actually reduced by the distractor, then the classification accuracy in this within-classification should be lower than in the original within-classification scheme where the classifier was trained and tested using the without-distractor trials. Moreover, in this cross-classification (i.e., trained on with-distractor and tested on without-distractor trials), we expect the classification accuracy to be higher than in the training on without-distractor and testing on with-distrctor trials cross-classification as the training will learn the more difficult comparison but then be tested on an easier set. Lower accuracy might suggest a transformation of information instead of reduced information.

In sum, we used four classification conditions in the main analysis: (1) train and test on without-distractor trials (within-classification), (2) train on without-distractor and test on with-distractor trials (cross-classification), (3) train and test on with-distractor trials (within-classification), and (4) train on with-distractor and test on without-distractor trials (cross-classification). To disclose the cortical region where imagery content is reduced by the distractor, we expected the classification accuracy to be (i) higher in (1) than in (2), (ii) higher in (1) than in (3), and (iii) higher in (4) than in (2).

## 3 Results

### 3.1 Behavioral task and vividness of visual imagery

When calculating the hit rates in the behavioral task during fMRI scanning (i.e., whether the red dot fell inside the area of the imagined shape) the whole area within the placeholders was considered the imagined shape. Two participants had chance level performance in the behavioral task and were thus excluded from further analyses. The average hit rate in the task was 89.6 % (SD 7.8 %). The hit rates between trials without distractor (mean±SD 91.4±6.3 %) and with distractor (mean±SD 87.8±9.9 %) were significantly different (paired t-test; t= 2.74, p=0.014, df=17, Cohen’s *d*=0.44), thus indicating that the distractor had a small effect on performance. No difference was found between the respective reaction times. All participants reported to have noticed the distractor consciously (at least on some trials as its visibility was not assessed on trial-by-trial basis). The mean score in the Vividness of visual imagery Questionnaire (VVIQ) was 3.48 (SD=0.68). One participant had the minimum score, indicating inability to form a visual image, and was therefore excluded from further analyses.

### 3.2 MVPA

The key analysis in this study relates to the question of in which brain area(s) does a concurrently presented visual distractor modulate imagery content representations; these results are shown in **Figure 3**, with the key findings of brain areas in which representations of imagery content are affected by visual distractor, in **Figure 3B** and **Table 2**.

**Figure 3:**
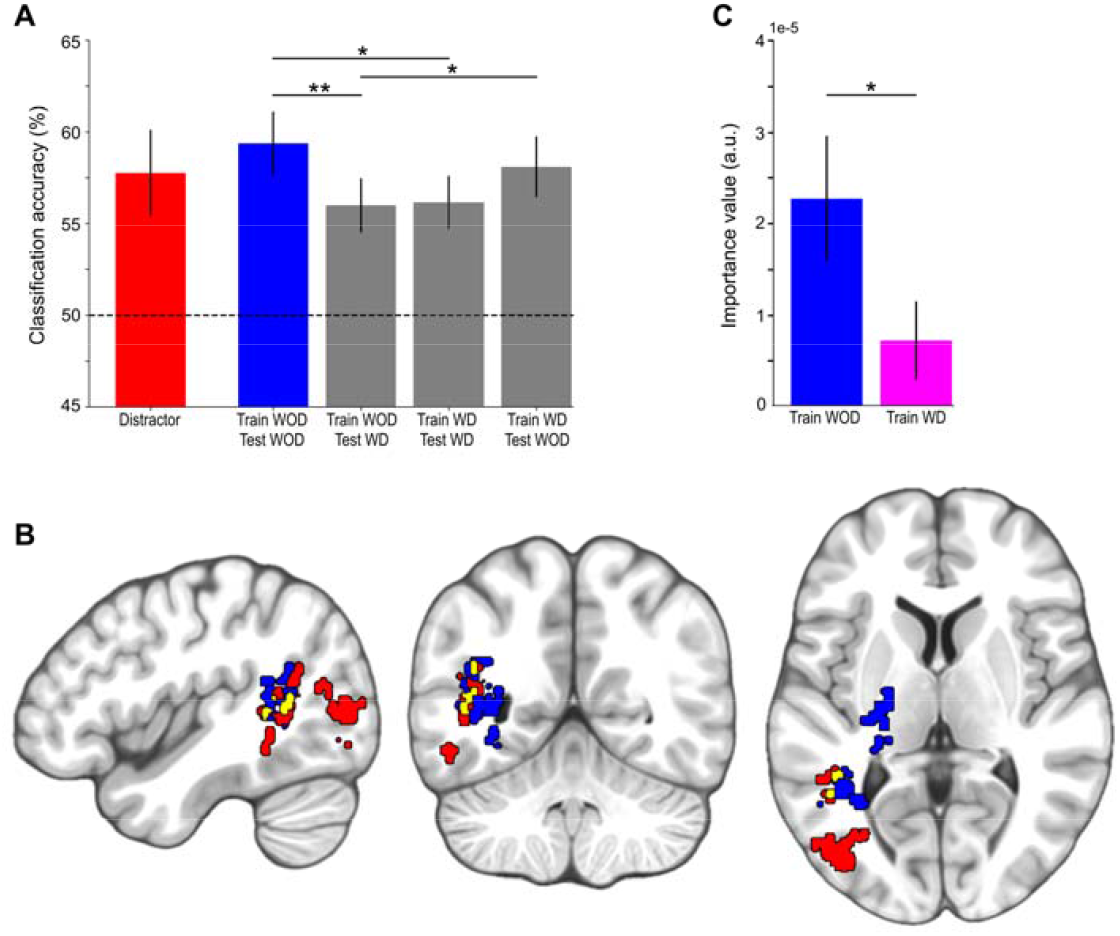
Classification accuracies and importance maps for the visual distractor and imagery conditions. **A)** Mean ± SEM classification accuracies for the classification conditions. The horizontal dashed line indicates the theoretical chance level accuracy (50%). All accuracies are statistically significant with permutation test p<0.001. Significant differences between the conditions are marked with asterisks (two-sample t-test: *p<0.05, **p<0.01). WOD: without distractor; WD: with distractor. **B)** Importance maps for the distractor (red) and imagery WOD (blue) classification conditions. For the distractor condition, the importance map for the with-distractor category is shown (see section 2.4). For the imagery WOD condition, the importance values for both categories (i.e., circle and diamond imagery) were combined into one imagery content representation map. The highest 5% of the importance values for each condition were retained and the remaining clusters larger than 125 (5×5×5) voxels are shown on sagittal (MNI x=-44), coronal (MNI y=-54), and axial (MNI z=6) slices (neurological convention, left is left). The areas in yellow depict the overlap of the importance maps between the two conditions. **C)** Mean ± SEM importance values for the imagery WOD and imagery WD classifiers within the regions that most contributed to both the perception of the distractor and the imagery of the shapes (i.e., areas depicted in yellow in B). For the importance maps for imagery WD classifier, see SI Fig. S1.

**Table 1** shows the classification accuracies for the distractor and imagery conditions (corresponding to **Fig. 3A**). All accuracies were significantly above chance level with permutation test p<0.001. The classification accuracy in the imagery condition where the classifier was trained and tested on without-distractor trials was significantly higher than the classification accuracies in the imagery conditions where the classifier was trained on without-distractor trials and tested on with-distractor trials (paired-samples t-test: t=3.2, p=0.005; **Fig. 3A**) and where the classifier was trained and tested on with-distractor trials (paired-samples t-test: t=2.1, p=0.025; **Fig. 3A**). Also, the classification accuracy in the the imagry condition where the classifier was trained on with-distractor trials and tested on without-distractor trials was higher than in the imagry conditions was trained on without-distractor trials and tested on with-distractor trials (paired-samples t-test: t=1.8, p=0.046; **Fig. 3A**). The importance values extracted from the brain regions that most contributed to both the perception of the distractor and the imagery of the shapes (i.e., the overlap depicted in yellow in **Fig. 3B**), were significantly decreased when the distractor was presented during the imagery (paired-samples t-test: t=2.2, p=0.02; **Fig. 3C**).

**Table 1:** Classification accuracies (mean ± SEM) in the distractor and imagery classification conditions. TrWOD: trained without distractor; TeWOD: tested without distractor; TrWD: trained with distractor; TeWD: tested with distractor.

| Condition | Classification accuracy (%) |
| --- | --- |
| Distractor | $57.8 \pm 2.3$ |
| Imagery (TrWOD, TeWOD) | $59.4 \pm 1.7$ |
| Imagery (TrWOD, TeWD) | $56.0 \pm 1.4$ |
| Imagery (TrWD, TeWD) | $56.1 \pm 1.4$ |
| Imagery (TrWD, TeWOD) | $58.1 \pm 1.6$ |

## 4 Discussion

The aim of this study was to determine the brain areas where visually presented information disrupts or weakens imagery representations. This was accomplished by using MVPA based on artificial neural networks on fMRI data collected using a behavioral paradigm in which participants were required to engage in imagery of visual shape while a distractor (in the form of visual shape) was presented on a proportion of trials. Strongest imagery representations were present in the left ventral temporal, lateral occipitotemporal, and inferior parietal cortices, involving the fusiform, lingual, posterior middle temporal, angular, and supramarginal gyri (see **Figure 3B and Table 2**). Additional areas implicated in imagery were the left insular and premotor cortices as well as brainstem. This finding is in line with previous fMRI studies that have been carried out on the topic (Albright, 2012; Ishai et al., 2000; Spagna, Hajhajate, Liu, & Bartolomeo, 2021; Winlove et al., 2018). Similarly, the presentation of the visual distractor was found to be most stongly represented in the lateral occipitotemporal and inferior parietal cortices, involving, for example, the middle occipital, posterior middle temporal, and angular gyri (see **Figure 3B** and **Table 2**).

**Table 2:** Clusters of highest importance values in the perception of the distractor, imagery without distractor (WOD), and imagery with distractor (WD) classification conditions (corresponding to red, blue, and magenta in Figure 3). The regions of overlap between the importance values in the perception and imagery conditions, corresponding to yellow and cyan colors in Figure 3, are also reported. Region label, number of voxels, and peak MNI coordinates are shown for each cluster.

|  | Region | Voxels | Peak MNI |  |  |
| --- | --- | --- | --- | --- | --- |
|  |  |  | x | y | z |
| Perception<br>(Distractor) | Middle occipital gyrus (L) | 169 | -42 | -78 | 6 |
|  | Angular gyrus (L) | 144 | -40 | -56 | 26 |
| Imagery WOD | Lingual gyrus (L) | 305 | -32 | -54 | 6 |
|  | Insula (L) | 226 | -32 | -20 | 14 |
|  | Precentral gyrus (L) | 217 | -20 | -8 | 52 |
|  | Brainstem | 186 | -10 | -14 | -18 |
| Imagery WD | Inferior parietal lobule (L) | 450 | -36 | -38 | 36 |
|  | Pallidum (R) | 285 | 22 | -18 | -2 |
|  | Pallidum (L) | 206 | -22 | -8 | 0 |
|  | Putamen (R) | 204 | 24 | 10 | 10 |
|  | Precentral gyrus (L) | 161 | -26 | -8 | 50 |
| Overlap of<br>perception and<br>imagery WOD | Middle temporal gyrus (L) | 17 | -44 | -52 | 6 |
|  | Angular gyrus (L) | 3 | -40 | -54 | 24 |
|  | Fusiform gyrus (L) | 2 | -36 | -44 | -10 |
| Overlap of perception and<br>imagery WD | Precentral gyrus (L) | 44 | -26 | -8 | 50 |
|  | Thalamus (L) | 16 | -24 | -32 | 12 |
|  | Middle temporal gyrus (L) | 14 | -32 | -54 | 8 |
|  | Insula (L) | 4 | -26 | -22 | 20 |
|  | Putamen (L) | 4 | -22 | -18 | 6 |
|  | Rolandic operculum (L) | 3 | -30 | -32 | 18 |

The novel aspect of the present study was to reveal the locus of imagery-vision interaction. As hypothesized, we found that the classification accuracy of the imagined shapes was significantly lower when using a classifier that was trained on without-distractor trials but tested on with-distractor trials compared to when the classifier was tested on without-distractor trials (see **Figure 3A**), implying that the distractor weakens the representation patterns underlying shape imagery. Similar decrease in classification accuracy was observed also when comparing the classifier that was trained and tested on without-distractor trials to a classifier that was trained and tested on with-distractor trials, acting as a control comparison as both classifiers were trained and tested using the same trial types (i.e., within-classification rather than cross-classification, see section 2.4). As a further control, we observed significantly higher classification accuracy for a classifier that was trained on with-distractor trials but tested on without-distractor trials as compared to a classifier that was trained on without-distractor trials but tested on with-distractor trials (i.e., cross-classification in both conditions).

Disclosing the brain areas that most contributed to the classifications, our main finding was that an area in the left lateral occipitotemporal cortex, centered in the occipitotemporal part of middle temporal gyrus and involving also the posterior part of fusiform gyrus and inferior part of angular gyrus, contains imagery content representations that were disrupted when the visual distractor was presented during imagery, indicating similarity of neural representations of imagery and perception (see **Table 2** and **Figure 3B)**. Importantly, as shown in **Figure 3C**, the contribution of these areas to the classification of shape imagery was significantly decreased when the distractor was presented (see also **SI Fig. S1**). The lateral occipitotemporal cortex is known to represent object and shape information (Radoslaw Martin Cichy, Chen, & Haynes, 2011; Eger, Ashburner, Haynes, Dolan, & Rees, 2008; Grill-Spector et al., 1999; Op de Beeck, Torfs, & Wagemans, 2008; Vinberg & Grill-Spector, 2008; Vuilleumier, Henson, Driver, & Dolan, 2002) and its involvement is likely to reflect the involvement of object-selective brain areas in both imagery maintenance and the encoding of distractor shape where their representations interact. In summary, our results support the hypothesis that imagery-vision interaction occurs relatively early on in the processing of incoming information rather than in higher-level regions.

Our study links well to prior work on distractor inference on visual short-term memory (VSTM) showing that distracting visual information affects VSTM representations in early visual areas but not in higher-level areas (Bettencourt & Xu, 2016; Lorenc et al., 2018). Given the close correspondence between neural representations of imagery and short-term memory (e.g., Albers et al., 2013), our findings can be easily reconciled with the evidence from short-term memory. To the best of our knowledge this is the first study to demonstrate the effect of distractors on visual imagery using MVPA.

### 4.1 Limitations

The findings of our study should be interpreted in the context of its limitations. First, a critical aspect of conscious perception is that the source of the percept needs to be ascertained (i.e., whether it is internally generated or externally induced) and the failure to do so will be detrimental to the ability to engage in the visual world (and for example, can be reflected as hallucinations). The Perky effect (1910), which reflects the intrusion and distraction of visual input into imagery, is one behavioral paradigm examining the process. It is important to note, however, that our paradigm differed from the Perky effect in that we did not assess distractor visibility and, as stated by the participants after the experiment, the distractor was visible.

Another possible limitation arises from the interpretation of the importance maps in terms of information representation in the brain (e.g., Haufe et al., 2014). That is, determining which voxels contribute most to the classification is not directly translatable to which voxels represent information about a given category. However, instead of extracting the raw weights of the classifier, whose magnitude and sign have no unambiguous interpretation, we used an importance extraction method based on layer-wise relevance propagation that decomposes the output of the classifier back to the inputs, thus providing a direct interpretation of each importance map (Bach et al., 2015; Montavon et al., 2017). Such importance extraction method for neural network classifier was recently shown to provide increased statistical power compared to univariate approaches for both detection of overlapping patterns as well as patterns with weak univariate information (Gotsopoulos et al., 2018).

In the main analysis, the importance maps of the imagery conditions were interpreted to contain imagery content representations. However, imagery content representations could be present in the voxel activation pattern even though no overall difference in activation is observed between conditions, and instead of containing imagery content representations, the activations could be related to some other aspect of the task or even be spurious (e.g., draining veins). Thus, some caution should be exercised when interpreting the importance maps as containing imagery content representations.

A statistically significant difference was observed in the hit rates between trials without and with distractor, implying that the distractor affected performance and therefore could constitute a confound in the main analysis. To alleviate this concern, the subjective difficulty of the task (defined as the difference in the hit rates between trials with and without distractor) was regressed out from the BOLD time series before the across-participants classification. As discussed above, the robustness of representations in temporo-parietal junction/supramarginal gyrus need to be interpreted in this context and could be taken to indicate that these representations more closely correlate with imagery task performance, given that they were not weakened by distractor presentation even when the subjective difficulty of the task was regressed out of the analysis.

### 4.2 Conclusions

Behaviorally, there is much evidence that internally generated and maintained visual representations are modulated by incoming visual information. As discussed in the Introduction, the principle was first demonstrated by Perky (1910) who demonstrated an intrusion of subliminal visual input into imagery representations. Subsequent studies have found similar effects in studies involving short-term memory as well (Bona, Cattaneo, Vecchi, Soto, & Silvanto, 2013; Lorenc et al., 2018; Magnussen, Greenlee, Asplund, & Dyrnes, 1991). These interactions have been shown to follow principles of visual cortical organization (e.g., masking effects follow principles of orientation channels; Magnussen et al, 1991; Bona et al, 2013). Our results provide a neural underpinning to these effects in the context of shape imagery. Specifically, when imagery and incoming visual input involve shape information, the interaction between the two occurs in object-selective lateral extrastriate cortex at the junction of left occipital, temporal and parietal lobes.

## Acknowledgements

We thank Marita Kattelus for her help in fMRI data collection. We acknowledge the computational resources provided by the Aalto Science-IT project.

## References

Albers, A. M., Kok, P., Toni, I., Dijkerman, H. C., & de Lange, F. P. (2013). Shared Representations for Working Memory and Mental Imagery in Early Visual Cortex. Current Biology, 23(15), 1427–1431. https://doi.org/https://doi.org/10.1016/j.cub.2013.05.065

Albright, T. D. (2012). On the Perception of Probable Things: Neural Substrates of Associative Memory, Imagery, and Perception. Neuron. https://doi.org/10.1016/j.neuron.2012.04.001

Bach, S., Binder, A., Montavon, G., Klauschen, F., Müller, K. R., & Samek, W. (2015). On pixel-wise explanations for non-linear classifier decisions by layer-wise relevance propagation. PLoS ONE, 10(7). https://doi.org/10.1371/journal.pone.0130140

Behrmann, M., Moscovitch, M., & Winocur, G. (1994). Intact Visual Imagery and Impaired Visual Perception in a Patient With Visual Agnosia. Journal of Experimental Psychology: Human Perception and Performance, 20(5), 1068–1087. https://doi.org/10.1037/0096-1523.20.5.1068

Bettencourt, K. C., & Xu, Y. (2016). Decoding the content of visual short-term memory under distraction in occipital and parietal areas. Nature Neuroscience, 19(1), 150–157. https://doi.org/10.1038/nn.4174

Bona, S., Cattaneo, Z., Vecchi, T., Soto, D., & Silvanto, J. (2013). Metacognition of visual short-term memory: Dissociation between objective and subjective components of VSTM. Frontiers in Psychology, 4(FEB). https://doi.org/10.3389/fpsyg.2013.00062

Cichy Radoslaw M., Heinzle, J., & Haynes, J. D. (2012). Imagery and perception share cortical representations of content and location. Cerebral Cortex, 22(2), 372–380. https://doi.org/10.1093/cercor/bhr106

Cichy Radoslaw Martin Chen, Y., & Haynes, J. D. (2011). Encoding the identity and location of objects in human LOC. NeuroImage, 54(3), 2297–2307. https://doi.org/10.1016/j.neuroimage.2010.09.044

Eger, E., Ashburner, J., Haynes, J. D., Dolan, R. J., & Rees, G. (2008). fMRI activity patterns in human LOC carry information about object exemplars within category. Journal of Cognitive Neuroscience, 20(2), 356–370. https://doi.org/10.1162/jocn.2008.20019

Farah, M. J., Levine, D. N., & Calvanio, R. (1988). A case study of mental imagery deficit. Brain and Cognition, 8(2), 147–164. https://doi.org/10.1016/0278-2626(88)90046-2

Gotsopoulos, A., Saarimäki, H., Glerean, E., Jääskeläinen, I. P., Sams, M., Nummenmaa, L., & Lampinen, J. (2018). Reproducibility of importance extraction methods in neural network based fMRI classification. NeuroImage, 181, 44–54. https://doi.org/10.1016/j.neuroimage.2018.06.076

Grill-Spector, K., Kushnir, T., Edelman, S., Avidan, G., Itzchak, Y., & Malach, R. (1999). Differential processing of objects under various viewing conditions in the human lateral occipital complex. Neuron, 24(1), 187–203. https://doi.org/10.1016/S0896-6273(00)80832-6

Haufe, S., Meinecke, F., Görgen, K., Dähne, S., Haynes, J. D., Blankertz, B., &Bießmann, F. (2014). On the interpretation of weight vectors of linear models in multivariate neuroimaging. NeuroImage, 87, 96–110. https://doi.org/10.1016/j.neuroimage.2013.10.067

Ishai, A., Ungerleider, L. G., & Haxby, J. V. (2000). Distributed neural systems for the generation of visual images. Neuron, 28(3), 979–990. https://doi.org/10.1016/S0896-6273(00)00168-9

Jacobs, C., Schwarzkopf, D. S., & Silvanto, J. (2018). Visual working memory performance in aphantasia. Cortex, 105, 61–73. https://doi.org/10.1016/j.cortex.2017.10.014

Keskar, N. S., Mudigere, D., Nocedal, J., Smelyanskiy, M., & Tang, P. T. P. (2016). On Large-Batch Training for Deep Learning: Generalization Gap and Sharp Minima. CoRR, abs/1609.0. Retrieved from http://arxiv.org/abs/1609.04836

Kosslyn, S. M. (2005). Mental images and the brain. Cognitive Neuropsychology. https://doi.org/10.1080/02643290442000130

Lee, S. H., Kravitz, D. J., & Baker, C. I. (2012). Disentangling visual imagery and perception of real-world objects. NeuroImage, 59(4), 4064–4073. https://doi.org/10.1016/j.neuroimage.2011.10.055

Lorenc, E. S., Sreenivasan, K. K., Nee, D. E., Vandenbroucke, A. R. E., &D’Esposito, M. (2018). Flexible coding of visual working memory representations during distraction. The Journal of Neuroscience, 3061–17. https://doi.org/10.1523/JNEUROSCI.3061-17.2018

Magnussen, S., Greenlee, M. W., Asplund, R., & Dyrnes, S. (1991). Stimulus-specific mechanisms of visual short-term memory. Vision Research, 31(7–8), 1213–1219. https://doi.org/10.1016/0042-6989(91)90046-8

Montavon, G., Lapuschkin, S., Binder, A., Samek, W., &Müller, K. R. (2017). Explaining nonlinear classification decisions with deep Taylor decomposition. Pattern Recognition, 65, 211–222. https://doi.org/10.1016/j.patcog.2016.11.008

O’Craven, K. M., & Kanwisher, N. (2000). Mental imagery of faces and places activates corresponding stimulus-specific brain regions. Journal of Cognitive Neuroscience, 12(6), 1013–1023. https://doi.org/10.1162/08989290051137549

Op de Beeck, H. P., Torfs, K., & Wagemans, J. (2008). Perceived Shape Similarity among Unfamiliar Objects and the Organization of the Human Object Vision Pathway. Journal of Neuroscience, 28(40), 10111–10123. https://doi.org/10.1523/JNEUROSCI.2511-08.2008

Pearson, J., Naselaris, T., Holmes, E. A., & Kosslyn, S. M. (2015). Mental Imagery: Functional Mechanisms and Clinical Applications. Trends in Cognitive Sciences. https://doi.org/10.1016/j.tics.2015.08.003

Perky, C. W. (1910). An Experimental Study of Imagination. The American Journal of Psychology, 21(3), 422. https://doi.org/10.2307/1413350

Reddy, L., Tsuchiya, N., & Serre, T. (2010). Reading the mind’s eye: Decoding category information during mental imagery. NeuroImage, 50(2), 818–825. https://doi.org/10.1016/j.neuroimage.2009.11.084

Spagna, A., Hajhajate, D., Liu, J., & Bartolomeo, P. (2021). Visual mental imagery engages the left fusiform gyrus, but not the early visual cortex: A meta-analysis of neuroimaging evidence. Neuroscience and Biobehavioral Reviews. https://doi.org/10.1016/j.neubiorev.2020.12.029

Vinberg, J., & Grill-Spector, K. (2008). Representation of Shapes, Edges, and Surfaces Across Multiple Cues in the Human Visual Cortex. Journal of Neurophysiology, 99(3), 1380–1393. https://doi.org/10.1152/jn.01223.2007

Vuilleumier, P., Henson, R. N., Driver, J., & Dolan, R. J. (2002). Multiple levels of visual object constancy revealed by event-related fMRI of repetition priming. Nature Neuroscience, 5(5), 491–499. https://doi.org/10.1038/nn839

Winlove, C. I. P., Milton, F., Ranson, J., Fulford, J., MacKisack, M., Macpherson, F., & Zeman, A. (2018). The neural correlates of visual imagery: A co-ordinate-based metaanalysis. Cortex, 105. https://doi.org/10.1016/j.cortex.2017.12.014

