## Supplementary material for "Where in the brain do internally generated and externally presented visual information interact?": SI

**Supplementary Information**

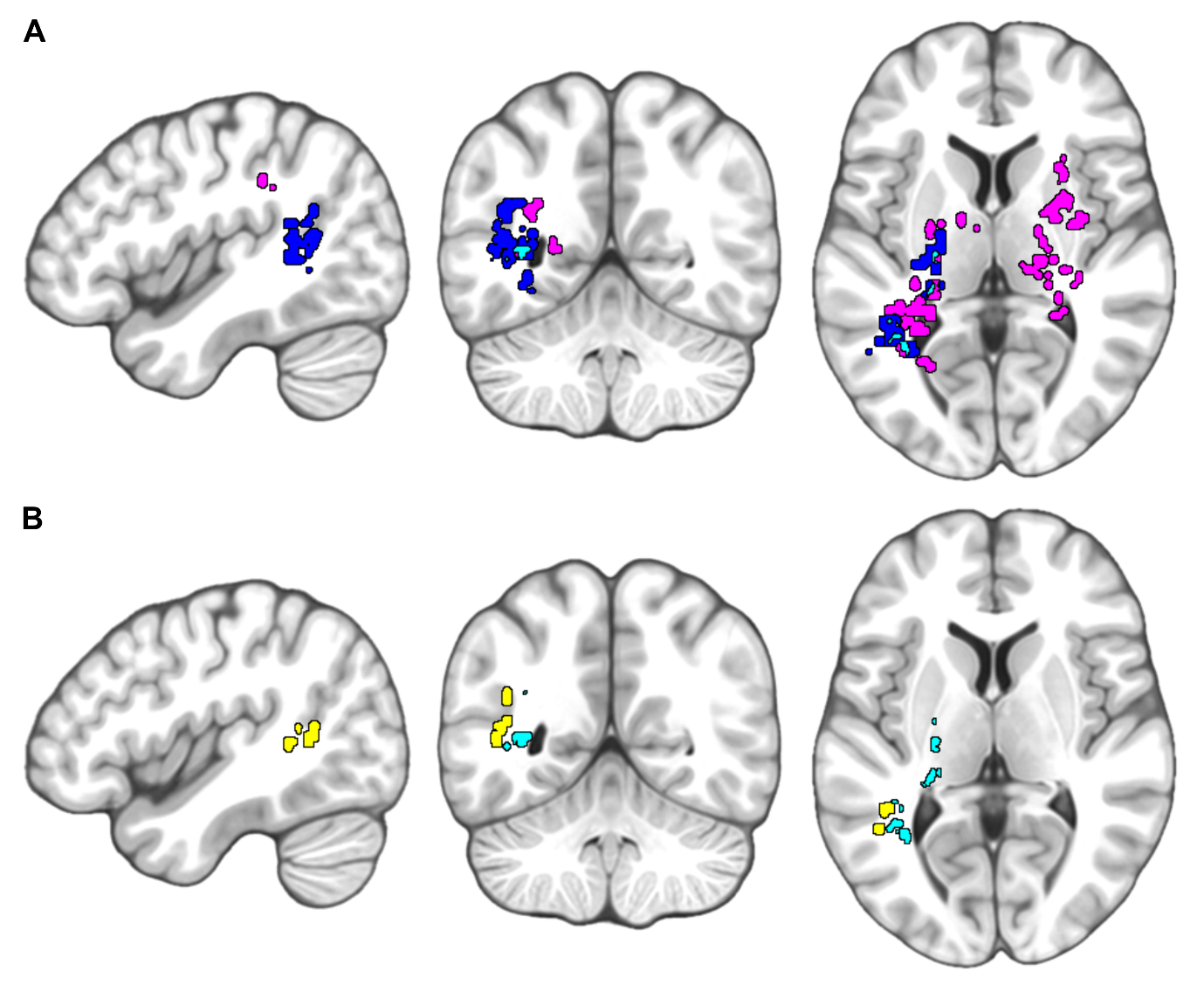

**Figure S1**: I**mportance maps for the imagery with and without distractor**. **A)** Importance maps for the imagery without distractor (blue) and imagery with distractor (magenta) classification conditions (importance values for circle and diamond imagery categories combined). The highest 5% of the importance values for each condition were retained and the remaining clusters larger than 125 (5×5×5) voxels are shown on sagittal (MNI x=-44), coronal (MNI y=-54), and axial (MNI z=6) slices (neurological convention, left is left). The areas in cyan depict the overlap of the importance maps between the two conditions. **B)** The overlap in A and the overlap of the importance maps for the distractor and imagery without distractor classification conditions (see Fig. 3B of the main text) plotted separately.
